# AI334 and AQ806 antibodies recognize the spike S protein from SARS-CoV-2 by ELISA

**DOI:** 10.1101/2020.05.08.084103

**Authors:** Philippe Hammel, Anna Marchetti, Wanessa C. Lima, Kelvin Lau, Florence Pojer, David Hacker, Pierre Cosson

## Abstract

We tested 10 recombinant antibodies directed against the spike S protein from SARS-CoV-1. Among them, antibodies AI334 and AQ806 detect by ELISA the spike S protein from SARS-CoV-2.

## Introduction

The spike (S) glycoprotein (UniProt #P0DTC2) mediates attachment of coronaviruses to the host ACE2 receptor (through the Receptor-Binding Domain [RBD] in the S1 subunit) and fusion with the host cell membrane (through the S2 subunit) (Yan *et al.*, 2020). Here we describe the ability of two recombinant antibodies (AI334 and AQ806) to detect by ELISA the soluble ectodomain of the S protein from SARS-CoV-2.

## Materials & Methods

### Antibodies

ABCD_AA831, ABCD_AF167, ABCD_AF618, ABCD_AH286, ABCD_AH287, ABCD_AH971, ABCD_AH974, ABCD_AI334, ABCD_AQ601, ABCD_AQ602, and ABCD_AQ806 antibodies (ABCD nomenclature, https://web.expasy.org/abcd/) were produced by the Geneva Antibody Facility (http://www.unige.ch/medecine/antibodies/) as mini-antibodies with the antigen-binding scFv portion fused to a mouse IgG2A Fc. The synthesized scFv sequences (GeneArt, Invitrogen) correspond to the sequences of the variable regions joined by a peptide linker (GGGGS)_3_ (see Table 1 for clone names and references). HEK293 suspension cells (growing in FreeStyle™ 293 Expression Medium, Gibco #12338) were transiently transfected with the vector coding for the scFv-Fc of each antibody. Supernatants (see Table 1 for individual yields) were collected after 5 days.

**Table 1:** Clone number, epitope, reference and production yields for the antibodies used in this study.

| ABCD | Clone | Epitope | Reference | Yield (mg/L) |
| --- | --- | --- | --- | --- |
| AA831 | 80R | S1/RBD | Sui *et al.*, 2004 | 100 |
| AF167 | S230.15 | S1/RBD | Rockx *et al.*, 2008 | 80 |
| AF618 | S227.14 | S1/RBD | Rockx *et al.*, 2008 | 80 |
| AH286 | m396 | S1/RBD | Prabakaran *et al.*, 2006 | 120 |
| AH287 | F26G19 | S1/RBD | Berry *et al.*, 2004 | <5 |
| AH974 | CR3013 | S1 | van den Brink *et al.*, 2005 | 120 |
| AI334 | CR3022 | S1 | ter Meulen *et al.*, 2006 | 50 |
| AQ601 | 3C7 | S1/RBD | Coughlin *et al.*, 2007 | 50 |
| AQ602 | 4D4 | S1 | Coughlin *et al.*, 2007 | <5 |
| AQ806 | VHH-72 | S1/RBD | Wrapp *et al.*, 2020a | 100 |
| AH971 | CR3009 | N protein | van den Brink *et al.*, 2005 | 150 |

### Antigen

The prefusion ectodomain (residues 1-1208) of the SARS-CoV-2 S protein, with a KV->PP substitution at residues 986/987, a RRAR->GSAS substitution at residues 682-685, and C-terminal T4 fibritin trimerization motif, protease cleavage site, TwinStrepTag and 8xHisTag (PDB #6VSB; Wrapp *et al.*, 2020b), was transiently transfected into 25×10^8^ suspension-adapted ExpiCHO cells (Thermo Fisher) using 1.5 mg plasmid DNA and 7.5 mg of PEI MAX (Polysciences) in 500 mL ProCHO5 medium (Lonza). Incubation with agitation was continued at 31°C and 4.5% CO_2_ for 5 days. The clarified supernatant was purified in two steps: via a Strep-Tactin XT column (IBA Lifesciences) followed by Superose 6 10/300 GL column (GE Healthcare) to a final concentration of 180 μg/ml in PBS.

### Protocol

S protein (10 μg/ml, 50 μl/well in PBS 0.5% (w/v) BSA, 0.1% (w/v) Tween20) was immobilized on streptavidin-coated ELISA plates (Pierce #15124) for 30 min. Each well was rinsed three times with 100 μl of washing buffer (PBS + 0.5% (w/v) BSA + 0.05% (w/v) Tween20), then incubated for 1 hour with 50 μl of each antibody-containing supernatant diluted in washing buffer (Fig. 1). After rinsing 3 times (100 μl washing buffer), wells were incubated with horseradish peroxidase-coupled goat anti-mouse IgG (Bio-Rad #170-6516, dilution 1:1000, 50 μl per well) for 30 min. After 3 rinses, Tetramethylbenzidine (TMB) substrate (Sigma #T5569) was added (50 μl per well). The reaction was stopped by the addition of 25 μl of 2 M H_2_SO_4_. The absorbance (OD) was measured at 450 nm, and the absorbance at 570 nm was subtracted.

**Fig. 1.**
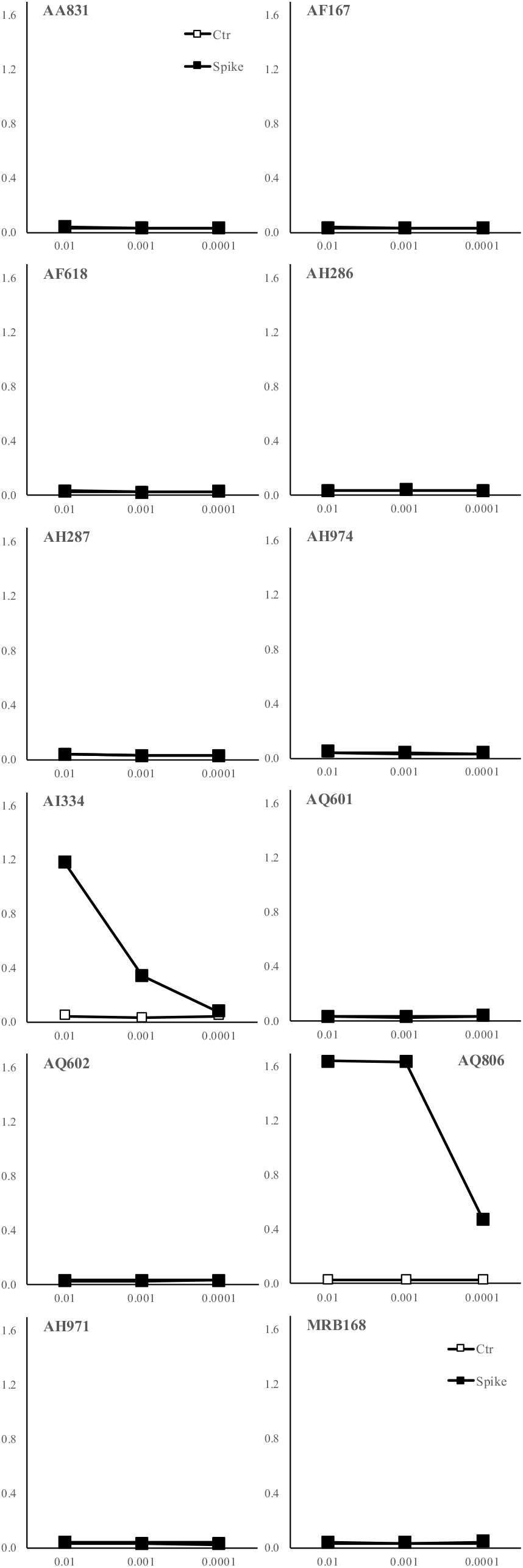
Specific binding of AI334 and AQ806 antibodies to the SARS-CoV-2 S protein, as detected by ELISA. On the Y axis, ELISA signal (in arbitrary units). On the X axis, the antibody dilution (1:100, 1:1’000 and 1:10’000). ‘Spike’ refers to the binding to the spike S protein; ‘Ctr’ refers to the binding to biotinylated BSA.

## Results

We tested 10 antibodies, originally developed against the SARS-CoV-1 S protein, for detection of the SARS-CoV-2 S protein by ELISA. From these, only two (AI334 and AQ806) bound in a concentration-dependent manner to the SARS-CoV-2 protein (Fig. 1). Two other antibodies, used as negative control (AH917 against the SARS-CoV-1 N protein and RB168 against an amoeba protein), also did not bind the S protein. AI334 and AQ806 antibodies have recently been shown to bind the RBD of the SARS-CoV-2 spike protein (Yuan *et al*., 2020; Wrapp *et al.*, 2020a).

## Acknowledgments

We would like to thank Prof. Jason McLellan (University of Texas, Austin) for providing the Spike expressing construct; and Laurence Durrer and Soraya Quinche (Protein Production and Structure Core Facility, EPFL) for the help with the mammalian cell culture.

## References

Berry JD, Jones S, Drebot MA, et al. Development and characterization of neutralizing monoclonal antibody to the SARS-coronavirus. J Virol Methods. 2004; 120:87–96. PMID:15234813

Coughlin M, Lou G, Martinez O, et al. Generation and characterization of human monoclonal neutralizing antibodies with distinct binding and sequence features against SARS coronavirus using XenoMouse. Virology. 2007; 361:93–102. PMID:17161858

Prabakaran P, Gan J, Feng Y, et al. Structure of severe acute respiratory syndrome coronavirus receptor-binding domain complexed with neutralizing antibody. J Biol Chem. 2006; 281:15829–36. PMID:16597622.

Rockx B, Corti D, Donaldson E, et al. Structural basis for potent cross-neutralizing human monoclonal antibody protection against lethal human and zoonotic severe acute respiratory syndrome coronavirus challenge. J Virol. 2008; 82:3220–35. PMID:18199635

Sui J, Li W, Murakami A, et al. Potent neutralization of severe acute respiratory syndrome (SARS) coronavirus by a human mAb to S1 protein that blocks receptor association. Proc Natl Acad Sci USA. 2004; 101:2536–41. PMID:14983044

ter Meulen J, van den Brink EN, Poon LL, et al. Human monoclonal antibody combination against SARS coronavirus: synergy and coverage of escape mutants. PLoS Med. 2006; 3:e237. PMID:16796401

van den Brink EN, Ter Meulen J, Cox F, et al. Molecular and biological characterization of human monoclonal antibodies binding to the spike and nucleocapsid proteins of severe acute respiratory syndrome coronavirus. J Virol. 2005; 79:1635–44. PMID:15650189

Wrapp D, De Vlieger D, Corbett KS, et al. Structural basis for potent neutralization of betacoronaviruses by single-domain camelid antibodies. Cell 2020a; pii:S0092-8674(20)30494-3. PMID:32375025

Wrapp D, Wang N, Corbett KS, et al. Cryo-EM structure of the 2019-nCoV spike in the prefusion conformation. Science. 2020b; 367:1260–1263. PMID:32075877

Yan R, Zhang Y, Li Y, Xia L, Guo Y, Zhou Q. Structural basis for the recognition of SARS-CoV-2 by full-length human ACE2. Science. 2020; 367:1444–1448. PMID:32132184

Yuan M, Wu NC, Zhu X, et al. A highly conserved cryptic epitope in the receptor-binding domains of SARS-CoV-2 and SARS-CoV. Science. 2020. pii:eabb7269. PMID: 32245784

